# Is there a neuropathic-like component to endometriosis-associated pain? Results from a large cohort questionnaire study

**DOI:** 10.1101/2020.07.01.181917

**Authors:** Lydia Coxon, Katja Wiech, Katy Vincent

## Abstract

**Background:** Pain is one of the primary symptoms of endometriosis, a chronic inflammatory condition characterised by the presence of endometrial tissue outside the uterus. Endometriosis-associated pain is commonly considered as nociceptive in nature but its clinical presentation suggests that it might have neuropathic-like properties in a subgroup of patients.

**Methods:** This is a cross sectional study using an online survey. The survey was distributed by patient support websites. The survey was composed of validated questionnaires assessing pain symptoms, psychological measures and questions about number of surgeries.

**Main results and the role of chance:** We had 1417 responses which met the inclusion criteria. Using standard painDETECT cut-off scores, we found that pain was classified as neuropathic in 40% of patients and as mixed neuropathic/nociceptive in a further 35%. In line with observations in other neuropathic conditions, the neuropathic subgroup reported higher pain intensities, greater psychological distress and cognitive impairment. Neuropathic pain was also more likely in those with more surgeries to the abdomen and a longer history of pain. As revealed by a cluster analysis, those with a neuropathic pain component could further be divided into two subgroups based on their sensory profile.

**Conclusions:** The data presented here indicate that endometriosis-associated pain includes a neuropathic-like component in a substantial proportion of women. Although further investigation is required, our finding challenges the current conceptualisation of endometriosis-associated pain as nociceptive and advocates for a new perspective on this type of pain, which is so debilitating to a large number of women.

## 1 Introduction

Endometriosis is a chronic inflammatory condition, characterised by the presence of endometrial tissue outside of the uterus^1^. Pain is a primary symptom, with dysmenorrhea, dyspareunia and non-cyclical pelvic pain being the most prevalent pain symptoms^2^. There is poor correlation between pain severity and disease burden and knowledge regarding biological mechanisms giving rise to pain is still sparse^3^. Currently available treatments focus on surgical excision/ablation of the lesions or hormonal suppression. Both of these are associated with risks and side effects, and have been linked to persistent or recurrent pain in a large proportion of women^4^.

In clinical practice, women often describe their endometriosis-related pain as ‘stabbing’ and ‘tingling’, which are known to be key features of neuropathic pain; however, the prevalence of a neuropathic-like component has not been properly investigated in this population. Neuropathic pain could be expected to arise in the context of endometriosis for a number of reasons. Firstly, there is currently no non-invasive test to establish the diagnosis of endometriosis. Consequently, all women with a confirmed diagnosis will have undergone at least one surgical procedure, with the associated risk of post-surgical neuropathic pain^5–7^. Secondly, endometriotic lesions are themselves innervated^2^. Surgical procedures excising/ablating the lesions may damage these nerve fibres, which could generate neuropathic pain^6^. Thirdly, these nerve fibres express TRPV1 receptors and are bathed in peritoneal fluid, known to contain high levels of inflammatory mediators such as BDNF and TNF-alpha in women with endometriosis, potentially sensitising nerve endings^2 8^. Thus, neuropathic pain could develop as a result of prolonged exposure to these inflammatory mediators^8^. In a recent study on chronic pelvic pain which included a subset of patients with endometriosis (n=32)^9^, over 50% of patients showed clinical features of neuropathic pain. There is ample evidence from both animal and human studies showing that neuropathic and nociceptive pain differ considerably, including with respect to their underlying pathology^10 11^, cognitive-affective processing^12 13^, and responses to treatment^10^. Therefore, a better understanding of the prevalence of neuropathic pain in endometriosis could guide both clinical care and future clinically oriented research strategies.

Here, we investigated the prevalence of neuropathic-like pain in endometriosis in a cohort of n=1417 women. To target a large sample from across the UK, we designed an online survey using painDETECT as a screening tool for neuropathic-like pain. We hypothesised that endometriosis-associated pain would be classified as neuropathic in a subgroup of women. As in other types of neuropathic pain^14^, we hypothesised that these women would score higher on questionnaires assessing depression and anxiety scores. We also expected the proportion of women with neuropathic-like pain to increase with the time since pain onset and with the number of surgeries to the abdomen. Studies in other neuropathic pain conditions have used cluster analysis to identify subtypes of neuropathic pain which differ in their sensory profile (using painDETECT responses)^115 16^. As these subtypes may have different responses to therapeutic agents, we used a similar strategy to test for subgroups amongst those within our cohort classified as having neuropathic or mixed pain.

## 2 Methods

### Survey design

The survey was created using Online Surveys (https://www.onlinesurveys.ac.uk) and was designed to be self-completed without professional guidance by combining single questions with validated questionnaires. Questions included patients’ ratings of the maximum pain intensity experienced over the last 12 months, for dysmenorrhea, dyspareunia, non-cyclical pain, dyschezia and dysuria using a Numerical Rating Scale (NRS) anchored at 0=no pain and 10=worst imaginable pain; duration of each type of pain since onset of symptoms (years) and number of surgeries received to the abdomen.

For somatosensory symptoms of neuropathic pain, painDETECT^17 18^ was used. This validated questionnaire consists of nine questions, which assess the severity, course, quality and nature of the patient’s pain. For example, the questions require the patient to rate the intensity of symptoms including spontaneous burning pain and pain evoked by light pressure, the pain course pattern, and to indicate whether their pain radiates to other parts of the body. Intensity ratings for symptoms are scored between 0 (ever), 1 (hardly noticed), 2 (slightly), 3 (moderately), 4 (strongly) or 5 (very strongly). The total painDETECT score ranges between -1 and 38. PainDETECT standard cut-off scores were used to group participants into those with nociceptive (≤12), mixed (13-18) and neuropathic-like (≥19) pain^17^. This categorisation has been shown to correspond to clinical diagnoses of neuropathic pain (e.g., neurological examination) in various chronic pain populations, including those not traditionally considered neuropathic^19^.

To assess a ‘centralised’ pain component, we included the Fibromyalgia Symptom Scale (FS)^20^ (scores are between 0 and 31) which assesses whether pain is widespread or localised and whether patients experience related symptoms. Although originally designed for fibromyalgia patients, the FS questionnaire has additionally been used in patients with postoperative pain following hysterectomy21 ^21^ as well as in other pain conditions ^22–24^.

For a specific assessment of symptoms of depression and anxiety we included the Beck Depression Inventory (BDI) (total score: 0-63)^25^; and State Trait Anxiety Inventory, Trait (STAI-T) (total score: 20-80)^26^.

Finally, the Pain Sensitivity Questionnaire (PSQ) was included as a patient-rated assessment of their pain sensitivity to hypothetical stimuli (0=not painful at all to 10=worst pain imaginable)^27^. PSQ ratings have been shown to be positively correlated with experimental pain intensity ratings and pain thresholds in healthy volunteers and chronic pain patients^27 28^.

The survey was posted on patient support websites (Endometriosis UK, www.endometriosis-uk.org; Endometriosis Association of Ireland, www.endometriosis.ie; and Endometriosis SHE Trust UK, www.facebook.com/EndoSheTrust/). It was open for responses between March and May 2018. Participants could complete the survey at their leisure, in several sessions and were able to withdraw at any time. The survey was beta tested on 9 patients, recruited from clinics at the Oxford University Hospitals Foundation Trust and the survey was modified according to their feedback. Patients were not reimbursed for their participation.

### Ethical approval

This study was approved by the Central University Research Ethics Committee, University of Oxford, R56567/RE002). Implied consent was attained via tick boxes to ensure the participant was over the age of 18 and that they agreed to take part in study.

### Data analysis

All statistical analysis was carried out using SPSS Statistics version 25. Participants were excluded if they had not received laparoscopic surgery (needed to diagnose endometriosis) or if they had not provided the age at which they received the diagnosis. The duration of pain symptoms was calculated as the difference between participants’ current age and the time-point they first experienced symptoms. Maximum duration of pain was defined as the longest duration for either dysmenorrhea, dyspareunia or non-cyclical pain.

To explore whether neuropathic pain is related to higher pain intensity, we compared pain intensity ratings between the three painDETECT groups separately for each pain type. To explore the relationship between painDETECT and (i) FS Score and (ii) PSQ Scores, correlation coefficients were calculated. BDI scores and STAI-T scores were compared between the neuropathic, mixed and nociceptive groups.

The number of surgeries to the abdomen was used as a grouping variable: group 1 had one surgery, group 2 had two surgeries, group 3 had three to four surgeries and group 4 had five or more surgeries. To test whether neuropathic-like pain was more prevalent in those with more surgeries, we compared the proportion of patients classified as having neuropathic pain between groups 1 to 4.

As most variables were not normally distributed (indicated by significant Kolmogorov-Smirnov tests), we chose to use non-parametric tests throughout. Group differences were explored using Kruskal-Wallis tests; significant results were followed up with Mann-Whitney U tests. Multiple comparison correction was carried out (Bonferroni correction). Correlation analyses were performed using Spearman’s correlations. Additionally, we calculated partial correlations controlling for pain intensity. Rho values are interpreted such that rho<0.2 is ‘very weak’; rho between 0.2 and 0.39 are ‘weak’; rho between 0.4 and 0.59 are ‘moderate’; rho between 0.6 and 0.79 are ‘strong’; and rho>0.8 are ‘very strong’.

#### Cluster analysis

Sensory symptom profiles (based on symptoms in painDETECT) of participants categorised as having mixed and neuropathic pain were used in two-step cluster analysis. To account for inter-individual differences in pain sensitivity, painDETECT scores for each of these symptoms were recalculated by subtracting the mean across all seven responses from each individual response as described in Baron et al., 2009^15^. Scores larger than zero thereby indicate a sensation that is more intense than the average individual symptom score.

Two-step cluster analysis with log-likelihood as distance measure was used with these continuous variables. Note that previous studies had employed a slightly different approach in their cluster analysis ^15 16^. We used a two-step cluster analysis with the optimal number of clusters determined using the Schwarz Bayesian Criterion. In addition to the number of clusters identified, we report the quality of the suggested solution based on the silhouette coefficient (ranging between -1 and 1), which jointly considers cluster cohesion and separation. Furthermore, we report the importance of each symptom for cluster formation (predictive importance).

Non-parametric Mann-Whitney U tests were used to assess differences between clusters with respect to pain and psychological measures.

## 3 Results

1615 responses were received, with 1417 meeting the inclusion criteria. Participant demographics can be seen in Table 1.

**Table 1.** Demographics of participants. Median values, range and interquartile range (IQR) of responses given as data not normally distributed.

|  | **Median (range) [IQR]** |
| --- | --- |
| **Age (years)** | 33 (18-59) [27-38] |
| **Dysmenorrhea Numerical Rating Scale Score (0-10)** | 8 (0-10) [7-9] |
| **Dyspareunia Numerical Rating Scale Score (0-10)** | 7 (0-10) [5-8] |
| **Non-cyclical Pain Numerical Rating Scale Score (0-10)** | 8 (0-10) [6-9] |
| **Duration of Pain Symptoms (years)** | 18 (1-47) [13-24] |
| **Number of surgeries to the abdomen** | 2 (1-23) [1-4] |

**Table 2.** Numerical Rating Scale scores, scores of cognitive and affective measures for participants in each painDETECT group (neuropathic, mixed and nociceptive). IQR= Interquartile range

|  | **Neuropathic**<br>**(median, IQR)** | **Mixed (median,**<br>**IQR)** | **Nociceptive**<br>**(median, IQR)** |
| --- | --- | --- | --- |
| **Dysmenorrhea** | 9 (8-10) | 8 (7-9) | 8 (7-9) |
| **Dyspareunia** | 7 (6-9) | 7 (5-8) | 6 (4-8) |
| **Non-cyclical pain** | 8 (7-10) | 8 (6-9) | 7 (5-8) |
| **Dyschezia** | 8 (6-9) | 7 (5-9) | 6 (4-8) |
| **Dysuria** | 6 (2-7) | 4 (0-6) | 0 (0-5) |
| **Depression (BDI)** | 28 (19-38) | 24 (16-34) | 16 (11-26) |
| **Anxiety (STAI-T)** | 55 (47-64) | 53 (45-61) | 48 (38-56) |
| **Fatigue** | 3 (2-3) | 2 (2-3) | 2 (2-3) |
| **Trouble thinking /**<br>**remembering** | 2 (1-3) | 2 (1-2) | 2 (1-2) |
| **Waking up feeling tired** | 3 (2-3) | 3 (2-3) | 2 (2-3) |

Of the 1417 participants, 40% (n=558) were categorised as having neuropathic pain according to their painDETECT scores. Pain was classified as mixed nociceptive/neuropathic in a further 35% (n=488) and as nociceptive in the remaining 25% (n=358).

To test whether the classification of pain as neuropathic was related to more intense pain, we compared intensity ratings for all relevant types of pain between the three groups. These analyses showed significant differences in NRS scores for dysmenorrhea (χ^2^(2)=91.710, p=6.088×10-20), dyspareunia (χ^2^(2)=80.191, p<0.001), non-cyclical pain (χ^2^(2)=96.884, p<0.001), dyschezia (χ^2^(2)=70.162, p<0.001) and dysuria (χ^2^(2)=117.853, p<0.001). For all pain types, scores were highest for the neuropathic group followed by the mixed group. Post hoc tests showed that all pairwise group comparisons reached statistical significance (p< 0.001) with the exception of non-cyclical pain for which the difference between nociceptive and mixed groups did not withstand multiple comparison correction (p=0.027 uncorrected).

Correlation analysis between painDETECT scores and FS and PSQ, showed a moderate correlation between painDETECT and FS scores (rho=0.441, p<0.001) and no significant correlation between painDETECT and PSQ.

Scores for depression and anxiety were significantly different between groups (χ^2^(2)=130.907, p<0.001; χ^2^(2)=67.389, p<0.001 respectively). Fatigue and trouble thinking/remembering were significantly different between groups (χ^2^(2)=85.219, p<0.001, χ^2^(2)=68.349, p<0.001 respectively). Additionally the neuropathic group was more likely to report waking up feeling tired (χ^2^(2)=80.588, p<0.001). Post hoc tests showed significant differences in all pairwise comparisons between groups, with the neuropathic group showing the strongest impairment followed by the mixed group (p<0.05 for all post hoc comparisons).

Correlation analyses revealed a significant positive correlation between painDETECT scores and each of the cognitive-affective variables, even when controlling for pain scores (Table 3).

**Table 3:** Correlations between painDETECT score and cognitive-affective variables. Standard correlations show Spearman’s correlation coefficients. Partial correlations control for Numerical Rating Scale scores for dysmenorrhea, dyspareunia and non-cyclical pain. * = p<0.001; BDI= Beck Depression Inventory, STAI-T= State-Trait Anxiety Inventory.

|  | BDI | STAI | fatigue | trouble<br>thinking/remembering | waking<br>up tired |
| --- | --- | --- | --- | --- | --- |
| **painDETECT**<br>**(standard**<br>**correlations)** | 0.33* | 0.24* | 0.26* | 0.24* | 0.26* |
| **painDETECT**<br>**(partial**<br>**correlations)** | 0.27* | 0.19* | 0.19* | 0.18* | 0.19* |

Comparing painDETECT scores between groups with increasing numbers of surgical procedures revealed significant group differences (χ^2^(3)=19.756, p<0.001) (see Figure 1). Post hoc tests confirmed that group 4 with five and more surgeries had a significantly higher painDETECT score than group 1 that had undergone one surgery (p<0.001).

**Figure 1:**
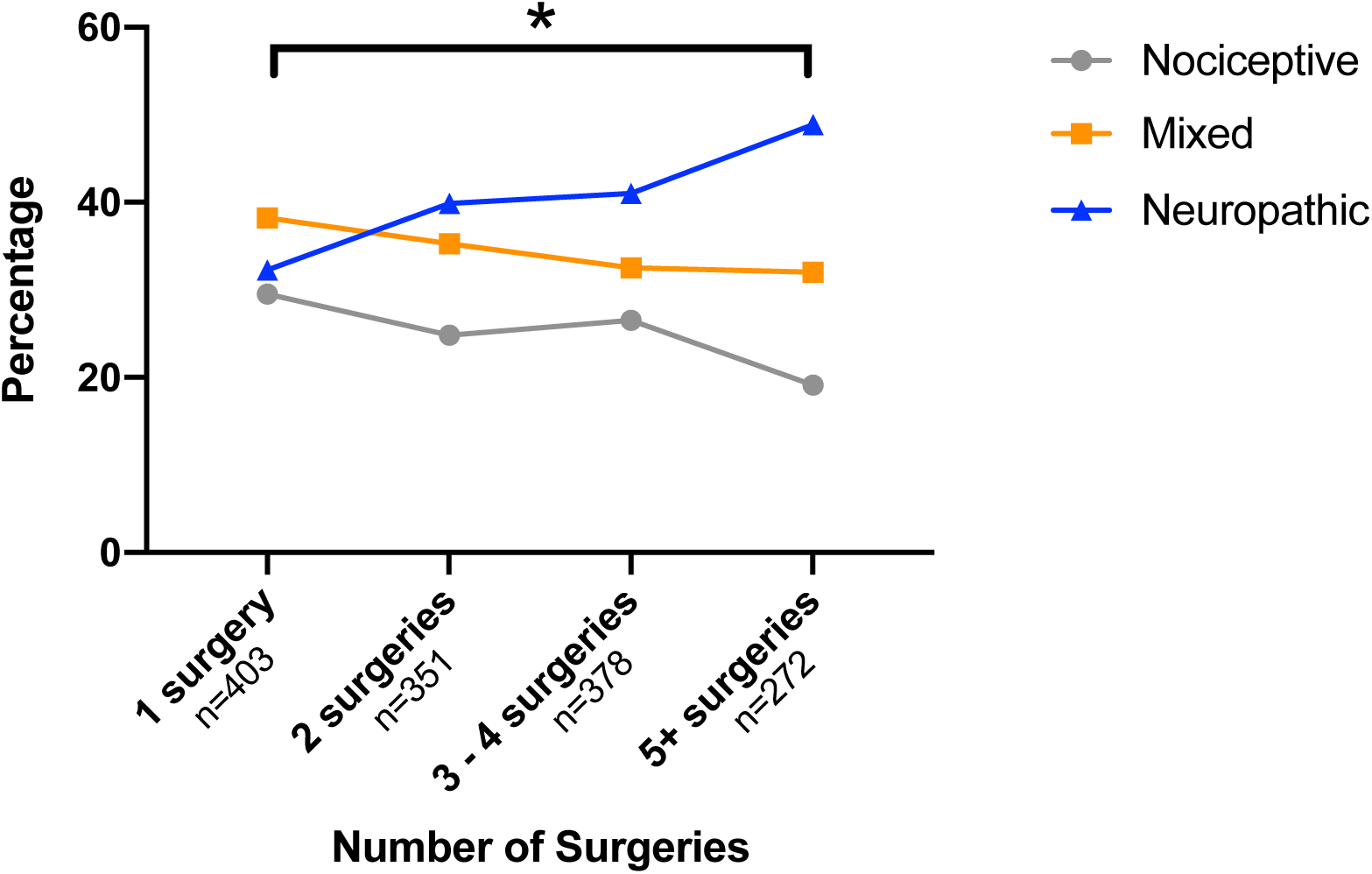
Relationship between painDETECT scores and number of abdominal surgeries. Surgeries were grouped so that each group had approximately the same number of participants. Surgery group 1 had one surgery to the abdomen, group 2 had two surgeries, group 3 had three to four surgeries and group 4 had five or more surgeries. Post hoc tests showed significant differences between group 1 and group 4 in painDETECT scores (p<0.001).

Exploring the relationship between duration of pain and painDETECT group, significant group differences were only found for duration of non-cyclical pain (χ^2^(2)=16.806, p<0.001). Post hoc tests showed that patients in the neuropathic group had experienced non-cyclical pain for longer than those in the mixed (p=0.006) and nociceptive (p<0.001) groups.

### Sensory symptom profiles and cluster analysis

To explore the sensory symptom profile of those categorised as having neuropathic or mixed pain, sensory symptoms rated in painDETECT were analysed in more detail. Table 4 shows the proportion of participants that had clinically relevant sensory disturbances (scores >3; strongly, very strongly). The presence of painful attacks was by far the most common symptom seen in this cohort.

**Table 4:** Reported symptoms in neuropathic and mixed groups. Proportion of participants in the neuropathic and mixed groups reporting clinically significant symptoms (i.e., a score of >3, strongly or very strongly) in the painDETECT questionnaire.

| Sensory symptom | Proportion of patients affected |
| --- | --- |
| Burning | 28.2% |
| Prickling | 25.4% |
| Mechanical Allodynia | 23.3% |
| Painful Attacks | 79.3% |
| Thermal Hyperalgesia | 8.9% |
| Numbness | 14.1% |
| Pressure Evoked Pain | 45.2% |

To determine if those categorised as having neuropathic or mixed pain could be divided into subgroups, we performed a cluster analysis based on their sensory symptom profiles. The analysis produced two clusters (Figure 2).

**Figure 2.**
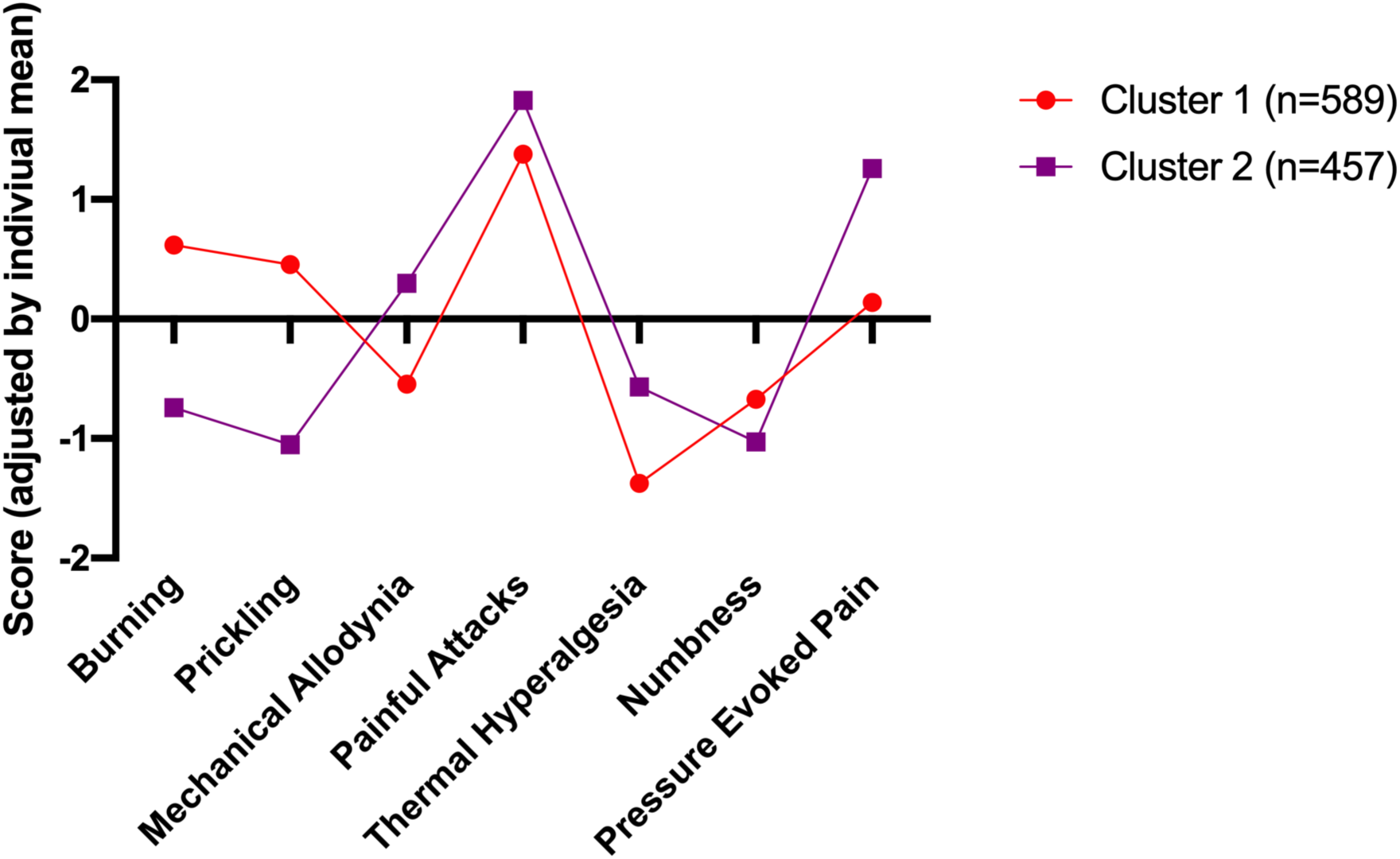
Sensory symptom profile of the two clusters. Scores were individually mean adjusted. The cluster model had a silhouette measure of cohesion and separation of 0.3, indicating it was of a ‘fair’ quality. 56.3% of patients fell into cluster 1 (n=589), and 43.7% in cluster 2 (n=457). The symptoms with the greatest predictive importance were ‘prickling/tingling’ (predictive importance=1), and ‘burning’ (predictive importance=0.77), which were most characteristic for cluster 1. In contrast, ‘pressure evoked pain’ (predictive importance=0.48), ‘thermal hyperalgesia’ (predictive importance=0.27) and ‘mechanical allodynia’ (predictive importance=0.27) were more common in cluster 2. ‘Painful attacks’ (predictive importance=0.11) and ‘numbness’ (predictive importance=0.05) were least discriminatory.

There were no significant differences between clusters with respect to depression (BDI), anxiety (STAI-T), fatigue, trouble thinking/remembering or waking up feeling tired (all p>0.05).

Clusters did not differ significantly with respect to the intensity of dysmenorrhea, dyspareunia or non-cyclical pain and PSQ scores (p>0.05) but FS scores were significantly higher in cluster 1 (p<0.001).

## 4 Discussion

Our data suggest that a considerable proportion of women with endometriosis-associated pain may have a neuropathic-like component to their pain. Based on painDETECT scores, pain was categorised as neuropathic in 40% of patients and as mixed (i.e., neuropathic and nociceptive pain) in a further 35% of the sample. In line with other chronic pain conditions where a mixture of underlying mechanisms, including neuropathic pain, is found^22 29 30^ women who were classified as having neuropathic-like pain also report higher pain intensity scores, more psychological distress and alterations in cognitive processing (Table 2). Interestingly, we found that women categorised as having neuropathic pain had undergone more surgical procedures and had a longer duration of non-cyclical pain than those in the other groups. Based on their sensory profile, those classified as having a neuropathic component could be further divided into two subgroups, suggesting that there may be more than one mechanism underlying neuropathic-like pain in women with endometriosis.

### Potential Mechanisms

As described, there are a number of potential mechanisms by which endometriosis may be associated with neuropathic-like pain. Here, we identified two factors related to the likelihood of neuropathic-like pain, namely the duration of non-cyclical pain and number of abdominal/pelvic surgeries (Figure 2). These relationships may simply represent failure of standard treatments when a neuropathic-like component is present, or they may in themselves be directly driving the neuropathic-like component. There is a growing body of evidence suggesting that surgery can generate neuropathic pain^5 7^, with it being more likely in women, those with pre-operative pain, psychological distress or inflammation^31 32^; all of which are frequently present in those undergoing surgery for endometriosis.

Our subsequent analyses do suggest that the variation in underlying mechanisms may be more limited in neuropathic-like pain associated with endometriosis than in other neuropathic conditions. For example, the majority (79%) of our participants reported painful attacks (as opposed to 32% in painful radiculopathy and 46% in postherpetic neuralgia)^16^. Furthermore, our cluster analysis (Figure 2) suggests that only two distinct subgroups of women exist.

### Clinical Relevance

Whilst endometriosis-associated pain syndrome is defined in the IASP taxonomy of pain^33^, clinically endometriosis is still predominantly managed by gynaecologists and the pain symptoms considered to arise either from the ectopic tissue implants themselves or the inflammatory environment of the pelvis. Thus, current guidelines^4 34 35^ all recommend simple analgesics (non-steroidal anti-inflammatories) and either hormonal suppression or surgical ablation/excision of the lesions. Pain management programmes are usually recommended only when treatment has failed. Many women with endometriosis-associated pain undergo repeated surgeries in the belief that pain symptoms may improve. A recent retrospective study found that over half the sample (n=486) had undergone >1 surgical procedure related to their endometriosis in a 10 year period^36^. All these interventions are based on the understanding that endometriosis-associated pain is nociceptive in nature. However, our data indicate, in line with other conditions associated with chronic pain (e.g. rheumatoid arthritis^30^, lower back pain^19 37^ and bladder pain syndrome^29^), a neuropathic-like component exists for a large proportion of women.

Our findings have direct implications for the treatment of endometriosis-associated pain. The high prevalence of neuropathic-like pain in our large sample clearly argues for a consideration of this type of pain, at the very least in those women with recurrent/persistent symptoms. The painDETECT questionnaire is a short and easy to complete screening tool used and validated in several chronic pain conditions^19^, which could be integrated into any standard gynaecology consultation. This is particularly true for those in whom a further surgical procedure is being considered. Whilst cross-sectional data such as this cannot determine underlying mechanisms or the direction of relationships, the strong relationship between the number of surgeries and a neuropathic-like component suggests that either surgery is not effective at treating this type of pain (leading to repeated procedures) or that it is in itself involved in generating this pain. Alternative strategies should therefore be considered first, unless there is another indication for surgery (e.g. pelvic mass). Furthermore, given the prevalence and associated personal and societal costs of endometriosis-associated pain^1^, attention should be given to determining effective medical strategies for treating neuropathic-like pain in these women. To date none of the available neuropathic adjunctive analgesics (e.g. amitriptyline, gabapentin etc) have been tested in endometriosis-associated pain, despite being relatively cheap and well-tolerated. Any such study should also take into account sensory symptom profiles.

### Limitations of the study

As participants were recruited from patient support groups, our sample might not necessarily be representative of the patient population. However, the long duration of pain and repeated surgical procedures we observed (Table 1) are commonly described for women with endometriosis^38 4 36^. Given that data were acquired using an online survey, we were unable to verify the diagnosis of endometriosis and we did not collect information on stage or location of disease for similar reasons. However, as there is known to be no correlation between stage/location and pain symptoms^2^ we do not consider that this would have influenced our overall findings.

painDETECT, since first publication in 2006^17^, has been used for several pain conditions, including those which are not normally thought of as neuropathic, such as rheumatoid arthritis, osteoarthritis, lower back pain, bladder pain and fibromyalgia in both the clinical and research setting^19^. Future longitudinal studies are needed to characterise underlying pathological processes and establish causality between these changes and the patients’ clinical presentation.

### Conclusion

The data presented here indicate that endometriosis-associated pain includes a neuropathic-like component in a substantial proportion of women. Our findings challenge the current conceptualisation of endometriosis-associated pain as nociceptive and advocates for a new perspective on this type of pain, which is so debilitating to a large number of women.

## Acknowledgements

Lydia Coxon has no conflicts of interest. Katja Wiech has received Consultancy fees from P&G Health, Germany. Katy Vincent declares that she has received research funding from Bayer AG, Honoraria from Eli Lilly and Honoraria and Consultancy fees from Bayer AG, Grunenthal GmBH and AbbeVie.

## Author Contributions

All authors have been involved in conception of project, collection of data, statistical analysis and preparation of manuscript.

## References

1. Zondervan KT, Becker CM, Koga K, Missmer SA, Taylor RN, Viganò P. Endometriosis. Nat Rev Dis Prim [Internet] 2018; 4: 9 Available from: https://doi.org/10.1038/s41572-018-0008-5

2. Coxon L, Horne AW, Vincent K. Pathophysiology of endometriosis-associated pain: A review of pelvic and central nervous system mechanisms. Best Pract. Res. Clin. Obstet. Gynaecol. 2018.

3. Vercellini P, Fedele L, Aimi G, Pietropaolo G, Consonni D, Crosignani PG. Association between endometriosis stage, lesion type, patient characteristics and severity of pelvic pain symptoms: A multivariate analysis of over 1000 patients. Hum Reprod [Internet] oup; 2007; 22: 266–71 Available from: http://dx.doi.org/10.1093/humrep/del339

4. Dunselman GAJ, Vermeulen N, Becker C, et al. ESHRE guideline: management of women with endometriosis †. Hum Reprod [Internet] 2014; 29: 400–12 Available from: https://doi.org/10.1093/humrep/det457

5. Lavand’homme P. Transition from acute to chronic pain after surgery. Pain [Internet] United States; 2017; 158: S50–4 Available from: http://insights.ovid.com/crossref?an=00006396-201704001-00007

6. Jones I, Bari F. Chronic pain after surgery. Surg [Internet] 2014; 32: 93–6 Available from: https://linkinghub.elsevier.com/retrieve/pii/S0263931913002640

7. Simanski CJP, Althaus A, Hoederath S, et al. Incidence of Chronic Postsurgical Pain (CPSP) after General Surgery. Pain Med (United States) 2014;

8. Sommer C, Leinders M, Üçeyler N. Inflammation in the pathophysiology of neuropathic pain. Pain 2018;

9. Whitaker LHR, Reid J, Choa A, et al. An Exploratory Study into Objective and Reported Characteristics of Neuropathic Pain in Women with Chronic Pelvic Pain. PLoS One [Internet] 2016; 11: e0151950. Available from: http://dx.doi.org/10.1371/journal.pone.0151950

10. St. John Smith E. Advances in understanding nociception and neuropathic pain. J. Neurol. 2018.

11. Colloca L, Ludman T, Bouhassira D, et al. Neuropathic pain. Nat Rev Dis Prim 2017;

12. Moriarty O, Ruane N, O’Gorman D, et al. Cognitive Impairment in Patients with Chronic Neuropathic or Radicular Pain: An Interaction of Pain and Age. Front Behav Neurosci 2017;

13. You Z, Zhang S, Shen S, et al. Cognitive impairment in a rat model of neuropathic pain: Role of hippocampal microtubule stability. Pain 2018;

14. Radat F, Margot-Duclot A, Attal N. Psychiatric co-morbidities in patients with chronic peripheral neuropathic pain: A multicentre cohort study. Eur J Pain (United Kingdom) 2013;

15. Baron R, Tölle TR, Gockel U, Brosz M, Freynhagen R. A cross-sectional cohort survey in 2100 patients with painful diabetic neuropathy and postherpetic neuralgia: Differences in demographic data and sensory symptoms. Pain 2009;

16. Mahn F, Hüllemann P, Gockel U, et al. Sensory symptom profiles and co-morbidities in painful radiculopathy. PLoS One 2011;

17. Freynhagen R, Baron R, Gockel U, Tölle TR. pain *DETECT* : a new screening questionnaire to identify neuropathic components in patients with back pain. Curr Med Res Opin 2006;

18. Mathieson S, Lin C. PainDETECT Questionnaire. J. Physiother. 2013. p. 211

19. Freynhagen R, Tölle TR, Gockel U, Baron R. The painDETECT project – far more than a screening tool on neuropathic pain. Curr Med Res Opin 2016;

20. Wolfe F, Clauw DJ, Fitzcharles MA, et al. Fibromyalgia criteria and severity scales for clinical and epidemiological studies: A modification of the ACR preliminary diagnostic criteria for fibromyalgia. J Rheumatol 2011; 38: 1113–22

21. Janda AM, As-Sanie S, Rajala B, et al. Fibromyalgia Survey Criteria Are Associated with Increased Postoperative Opioid Consumption in Women Undergoing Hysterectomy. Anesthesiology 2015;

22. Ramjeeawon A, Choy E. Neuropathic-like pain in psoriatic arthritis: evidence of abnormal pain processing. Clin Rheumatol 2019;

23. Kowalik CG, Cohn JA, Delpe S, et al. Painful Bladder Symptoms Related to Somatic Syndromes in a Convenience Sample of Community Women with Overactive Bladder Symptoms. J Urol 2018;

24. McKernan LC, Johnson BN, Crofford LJ, Lumley MA, Bruehl S, Cheavens JS. Posttraumatic Stress Symptoms Mediate the Effects of Trauma Exposure on Clinical Indicators of Central Sensitization in Patients with Chronic Pain. Clin J Pain 2019;

25. Dozois DJA. Beck Depression Inventory-II. Corsini Encycl Psychol 2010.

26. Spielberger R.; Lushere, R. C. G. State-Trait Anxiety Inventory. Prof Psychol 1971;

27. Ruscheweyh R, Marziniak M, Stumpenhorst F, Reinholz J, Knecht S. Pain sensitivity can be assessed by self-rating: Development and validation of the Pain Sensitivity Questionnaire. Pain 2009;

28. Ruscheweyh R, Verneuer B, Dany K, et al. Validation of the Pain Sensitivity Questionnaire in chronic pain patients. Pain 2012; 153: 1210–8

29. Cory L, Harvie HS, Northington G, Malykhina A, Whitmore K, Arya L. Association of neuropathic pain with bladder, bowel and catastrophizing symptoms in women with bladder pain syndrome. J Urol 2012;

30. Koop SMW, ten Klooster PM, Vonkeman HE, Steunebrink LMM, van de Laar MAFJ. Neuropathic-like pain features and cross-sectional associations in rheumatoid arthritis. Arthritis Res Ther 2015;

31. Zheng H, Schnabel A, Yahiaoui-Doktor M, et al. Age and preoperative pain are major confounders for sex differences in postoperative pain outcome: A prospective database analysis. PLoS One 2017;

32. Suffeda A, Meissner W, Rosendahl J, Guntinas-Lichius O. Influence of depression, catastrophizing, anxiety, and resilience on postoperative pain at the first day after otolaryngological surgery: A prospective single center cohort observational study. Med (United States) 2016;

33. Baranowski A, Abrams P, Berger RE, Buffington A, Collett B, Emmanuel A, Fall M, Hanno P, Howard FM HJ. Taxonomy of Pelvic Pain. Classification of Chronic Pain. IASP [Internet] 2012; Available from: https://www.iasp-pain.org/PublicationsNews/Content.aspx?ItemNumber=1673

34. National Institute for Health and Care Excellence. Guideline scope Endometriosis: diagnosis and management. Guidel Scope 2015;

35. Johnson NP, Hummelshoj L. Consensus on current management of endometriosis. Hum Reprod 2013;

36. Cheong Y, Tay P, Luk F, Gan HC, Li TC, Cooke I. Laparoscopic surgery for endometriosis: How often do we need to re-operate? J Obstet Gynaecol (Lahore) 2008;

37. Baron R, Binder A, Attal N, Casale R, Dickenson AH, Treede RD. Neuropathic low back pain in clinical practice. Eur. J. Pain (United Kingdom). 2016.

38. Nnoaham KE, Hummelshoj L, Webster P, et al. Impact of endometriosis on quality of life and work productivity: A multicenter study across ten countries. Fertil Steril 2011;

