## Supplementary Material for "Is there a neuropathic-like component to endometriosis-associated pain? Results from a large cohort questionnaire study"

### Supplementary Material – Hierarchical cluster analysis

#### Methods:

To ensure comparability with previous findings, our analysis pipeline followed the approach by Baron et al., 2009[15] who used a hierarchical WARD-approach with a squared Euclidian distance measure and a cut-off point of five clusters. We ran cluster analysis on both the neuropathic and mixed groups and neuropathic group alone. We set the number of clusters to 5 for each, as Baron et al. 2009[15].

#### Results:

For analysis of those in neuropathic and mixed groups 20% fell into cluster 1 (n= 212), 24% in cluster 2 (n= 247), 13% into cluster 3 (n= 132), 28% into cluster 4 (n=298) and 15% into cluster 5 (n= 157). The differences in sensory symptom profiles between the clusters can be seen in Supplementary Figure 1.

For analysis of those in the neuropathic group only 16% fell into cluster 1 (n= 88), 28% into cluster 2 (n= 154), 16% into cluster 3 (n= 92), 11% into cluster 4 (n= 59) and 30% into cluster 5 (n= 165). The differences in the sensory symptom profiles between clusters can be seen in Supplementary Figure 2.

#### Discussion:

This analysis was carried out to ensure comparability with previous studies, however in this setting we felt it was not appropriate so chose to report the two-step method. The two-step cluster method automatically selects the number of clusters based on the data which provides a more objective output. We also considered that forcing 5 clusters on our data did not give us clinically significant results and that fewer clusters were more meaningful in this setting.

Following the same method of cluster analysis did not give us clusters which were more comparable to those found in previous work, this is in part due to the fact that pain attacks are far more frequent in our population to those previously studied in this way.

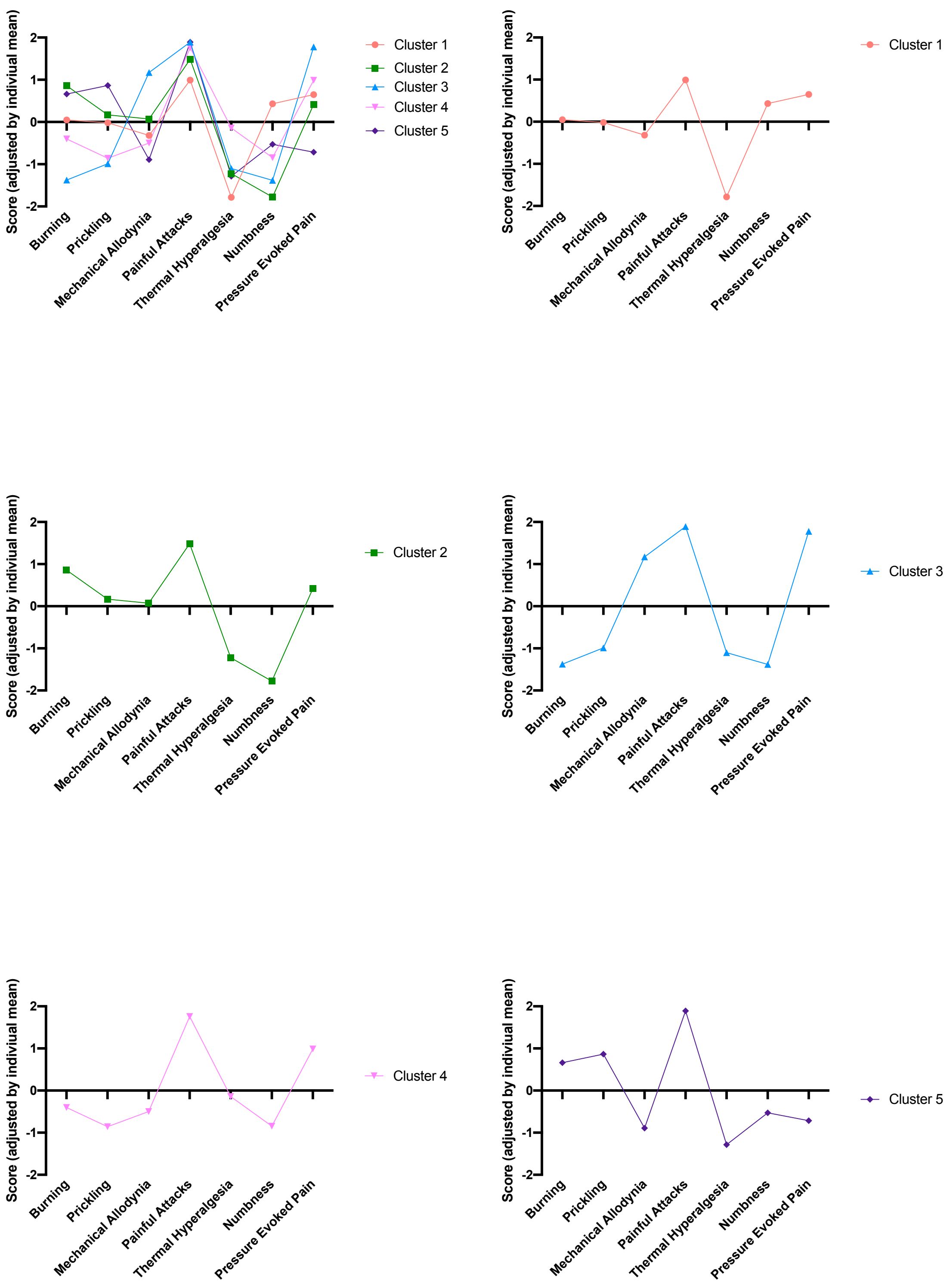


**Supplementary Figure 1. Sensory symptom profile comparison between clusters.** Sensory variables derived from painDETECT responses and were individually mean adjusted. Hierarchical cluster to create 5 clusters within neuropathic and mixed groups. This figure shows the symptom profile based on individually mean adjusted responses to painDETECT questions for the 5 clusters.

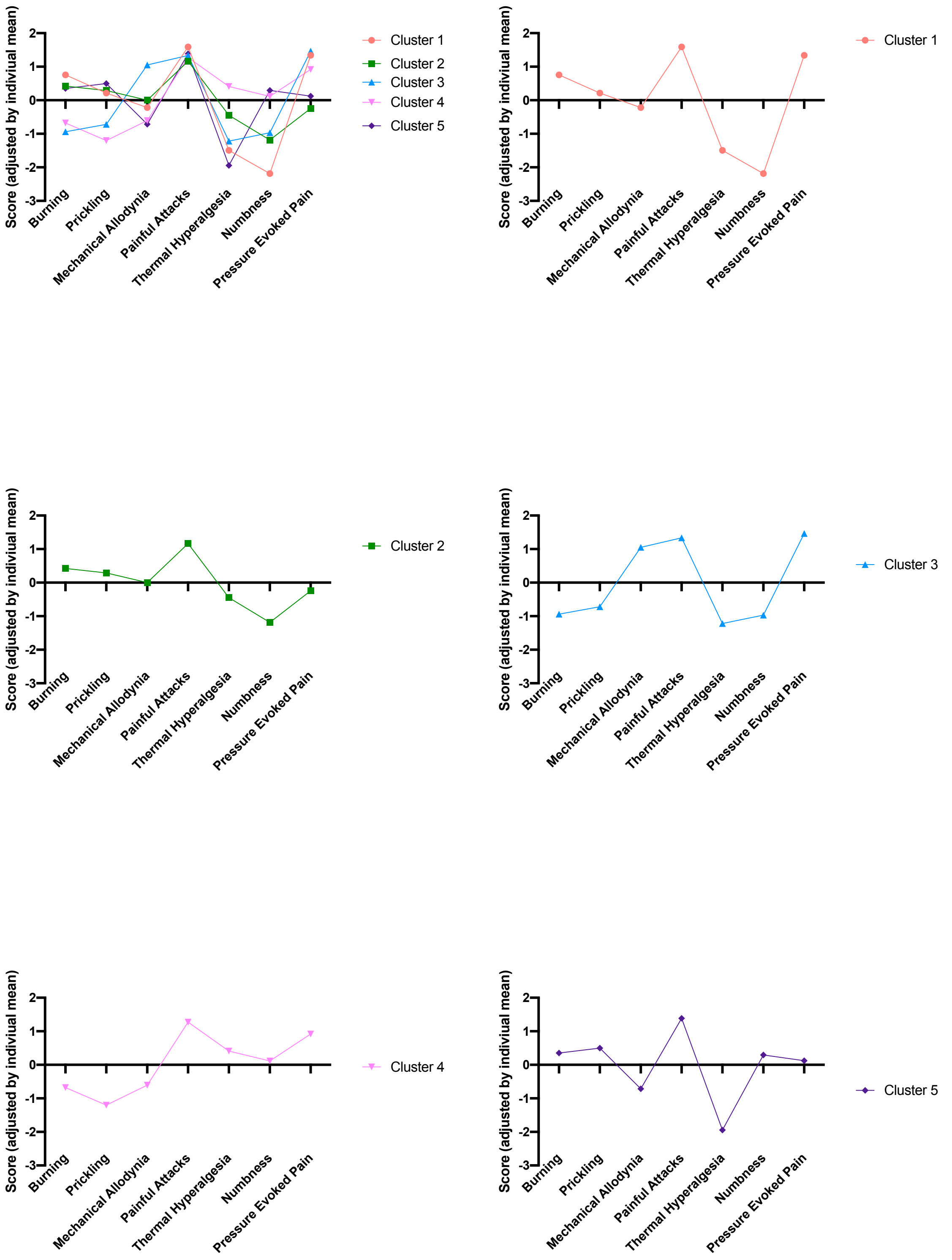


**Supplementary Figure 2. Sensory symptom profile comparison between clusters in neuropathic group.** Sensory variables derived from painDETECT responses and were individually mean adjusted. Hierarchical cluster to create 5 clusters within neuropathic group only. This figure shows the symptom profile based on individually mean adjusted responses to painDETECT questions for the 5 clusters.
